# Benchmarking optimization methods for parameter estimation in large kinetic models

**DOI:** 10.1101/295006

**Authors:** Alejandro F. Villaverde, Fabian Fröhlich, Daniel Weindl, Jan Hasenauer, Julio R. Banga

## Abstract

**Motivation:** Mechanistic kinetic models usually contain unknown parameters, which need to be estimated by optimizing the fit of the model to experimental data. This task can be computationally challenging due to the presence of local optima and ill-conditioning. While a variety of optimization methods have been suggested to surmount these issues, it is not obvious how to choose the best one for a given problem *a priori*, since many factors can influence their performance. A systematic comparison of methods that are suited to parameter estimation problems of sizes ranging from tens to hundreds of optimization variables is currently missing, and smaller studies indeed provided contradictory findings.

**Results:** Here, we use a collection of benchmark problems to evaluate the performance of two families of optimization methods: (i) a multi-start of deterministic local searches; and (ii) a hybrid metaheuristic combining stochastic global search with deterministic local searches. A fair comparison is ensured through a collaborative evaluation, involving researchers applying each method on a daily basis, and a consideration of multiple performance metrics capturing the trade-off between computational efficiency and robustness. Our results show that, thanks to recent advances in the calculation of parametric sensitivities, a multi-start of gradient-based local methods is often a successful strategy, but a better performance can be obtained with a hybrid metaheuristic. The best performer is a combination of a global scatter search metaheuristic with an interior point local method, provided with gradients estimated with adjoint-based sensitivities. We provide an implementation of this novel method in an open-source software toolbox to render it available to the scientific community.

**Availability and Implementation:** The code to reproduce the results is available at Zenodo https://doi.org/10.5281/zenodo.1160343

## 1 Introduction

Mechanistic kinetic models provide a basis to answering biological questions via mathematical analysis. Dynamical systems theory can be used to interrogate these kinetic models, enabling a more systematic analysis, explanation and understanding of complex biochemical pathways. Ultimately, the goal is the model-based prediction of cellular functions under new experimental conditions [1, 32, 34, 53]. During the last decade, many efforts have been devoted to developing increasingly detailed and, therefore, larger systems biology models [29, 49, 51]. Such models are often formulated as nonlinear ordinary differential equations (ODEs) with unknown parameters. As it is impossible to measure all parameters directly, parameter estimation (i.e. model calibration) is crucial for the development of quantitative models. The unknown parameters are typically estimated by solving a mathematical optimization problem which minimizes the mismatch between model predictions and measured data [2, 5, 28, 46].

Parameter estimation for dynamical systems is an inverse problem [55] that exhibits many possible challenges and pitfalls, mostly associated with ill-conditioning and non-convexity [48]. These properties, which are in general only known a posteriori, influence the performance of optimization methods. Even if we restrict our attention to a specific class of problems within the same field (e.g., parameter estimation in systems biology), there are often large differences in performance between different applications [31]. Hence, methods need to be benchmarked for a representative collection of problems of interest in order to reach meaningful conclusions. In this study, we consider the class of medium to large scale kinetic models. These models pose several challenges, such as computational complexity, and an assessment of the performance of optimization methods is particularly important [4, 13, 58].

The calibration of large-scale kinetic models usually requires the optimization of a multi-modal objective function [8,35,40], i.e. there will be several local optima. Local optimization methods, such as Levenberg-Marquardt or Gauss-Newton [48], which converge to local optima, will only find a global optimum for appropriate starting points. Convergence to a suboptimal solution is an estimation artifact that can lead to wrong conclusions: we might think that the mechanism considered is not suitable to explain the data, while the real reason might be that the method failed to locate the global optimum [6].

In order to avoid suboptimal solutions, many studies have recommended the use of global optimization techniques [3,5,8,37]. One of the earliest and simplest global optimization methods is the multi-start, which consists of launching many local searches from different initial points in parameter space, assuming that one of them will be inside the basin of attraction of the global solution. It has been shown that multi-starts of local optimization methods can be sufficient for successful parameter estimation in kinetic models [20, 46], although the use of other approaches, such as metaheuristics, has also been advocated [22, 58].

In this study, we evaluate the state-of-the-art in parameter estimation methodologies and provide guidelines for their application to large kinetic models in systems biology. To this end, we use seven previously published estimation problems to benchmark a number of optimization methods. The selected problems are representative of the medium and large scale kinetic models used in systems biology, with sizes ranging from dozens to hundreds of state variables and parameters (see Table 2 for details). To the best of our knowledge, this is the first time that a systematic evaluation of parameter estimation methods is conducted on a set of problems of this size and characteristics. We compare several variants of state-of-the-art optimization methods which have been recently reported as competitive options for large problems, including multi-start [46] and hybrid metaheuristics [58]. We perform systematic comparisons between these different approaches using metrics capturing the performance/robustness trade-offs. Finally, we discuss the implications of our results and provide guidelines for the successful application of optimization methods in computational systems biology.

## 2 Methods and Benchmark Problems

### 2.1 Problem definition: Parameter optimization for ODE models describing biological processes

We consider deterministic dynamic systems described by nonlinear ODEs of the following form:

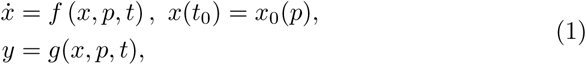

in which *x*(*t*) is the vector of state variables at time *t*, *x*_0_ is the vector of initial conditions, *f* is the vector field of the ODE, *g* is the observation function, and *p* is the vector of unknown constant parameters with lower and upper bounds *p^L^≤p≤p^U^*.

Parameter optimization for dynamical systems is a nonlinear dynamic optimization problem that aims to find the vector of parameter values *p* that minimizes the distance between model simulation and measured data subject to the dynamics of the system and (potentially) other possible constraints. The distance is measured by a scalar objective function (or cost function), which can be of several forms. One common choice is the weighted least squares objective function given by:

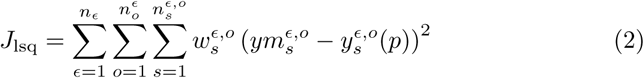

in which *n_E_* is the number of experiments, 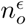 is the number of observables per experiment, 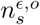 is the number of samples per observable per experiment, 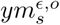 is the measured data, 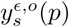 is the corresponding simulated output, and 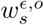 are constants that weight the observables in the objective function according to their magnitudes and/or the confidence in the measurements.

Another common choice for the objective function is the log-likelihood. Assuming independent, normally distributed additive measurement noise with standard deviation 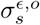, the likelihood of observing the data *D* given the parameters *p* is:

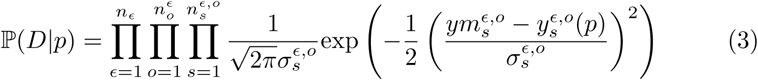

Maximizing (3) is equivalent to minimizing the negative log-likelihood function:

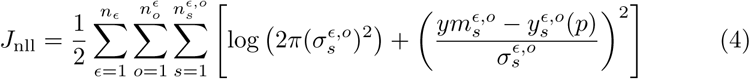

If the standard deviations 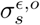 are known, *J*_lsq_ and *J*_nll_ possess the same optimal parameters. Furthermore, for 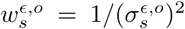, the log-likelihood and least squares functions are related by

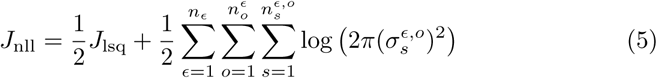

We remark that a good agreement of model output and data does not imply that the parameter estimates are correct or reliable. Practical and structural non-identifiabilities can prevent a parameter from being precisely determined [15]. Still, an accurate fit – and hence optimization – is the starting point for many uncertainty analysis methods. State-of-the-art identifiability analysis methods have been recently evaluated elsewhere [10, 33, 38, 45, 54].

### 2.2 Overview of optimization methods

The ideal optimization method for the above class of problems would be able to find the global optimum with guarantees and in a short computation time. Furthermore, it should scale well with problem size and be able to handle arbitrary non-linearities. Currently, no such method exists.

Local gradient-based methods [48] can be efficient but will converge to the local optimum in the basin of attraction where they are initialized. Local gradient-free (also called zero-order) methods, such as pattern search [59], are less efficient than gradient-based methods but more robust with respect to situations where the gradient is unavailable or unreliable [11].

Global methods aim to locate the global solution by means of either deterministic [19] or stochastic [60] strategies. Deterministic methods include so-called complete and rigorous approaches, both of which can ensure convergence to the global solution under certain circumstances. In contrast, stochastic (also known as probabilistic) methods can only guarantee global optimality asymptotically in the best case [42], but can solve many problems that cannot be handled using available deterministic methods. Both deterministic and stochastic global optimization methods have been used to solve parameter estimation problems in systems biology. The results show that deterministic methods suffer from lack of scalability [39]. The computational cost of purely stochastic methods (such as simulated annealing, or genetic algorithms) usually scales up better, but the computation times can still be excessive for problems of realistic size [37, 40].

Hybrid global-local methods attempt to exploit the benefits of global and local methods. By combining diversification phases (global search) and intensification phases (local search), hybrid methods facilitate reliable global exploration and fast local convergence. As a result, hybrid methods can potentially outperform the efficiency (convergence rate) of purely stochastic methods while keeping their success rate. One such hybrid method is the so called multi-start (MS) strategy [60], which solves the problem repeatedly with local methods initialized from different (e.g. random) initial points. Thus, MS can be regarded as one of the earliest hybrid strategies, and there are different extensions available [25, 60]. An alternative family of hybrid methods are metaheuristics (i.e. guided heuristics). An example is the enhanced scatter search (eSS) method [17], an improvement of the method designed by [23]. The eSS method combines a global stochastic search phase with local searches launched at selected times during the optimization, in order to accelerate convergence to local optima. Further accelerations can be achieved by parallelization [44, 57].

In all hybrid methods the efficiency of local methods plays a major role. The most efficient local methods are gradient-based, so their performance depends crucially on the accuracy of the gradient calculations [43]. The simplest way of approximating the gradient is by finite differences. However, more accurate gradients are provided by forward sensitivity analysis [47] and adjoint sensitivity analysis [20]. While the former provides information on individual residuals which can be used in least squares algorithms, the latter is more scalable.

### 2.3 Choice of optimization methods for benchmarking

In this study, we consider several competitive hybrid methods based on the recent results reported by [21] and [58]. These methods are summarized in Table 1 and combine two global strategies:

**Table 1:** Classification of the optimization methods considered in the benchmarking. These methods result from the combination of two global strategies with three local methods and two types of scaling for the search space.

| Global strategy | Local method & gradient calculation |  |  |  | Parameter scaling |
| --- | --- | --- | --- | --- | --- |
|  | FMINCON-ADJ | NL2SOL-FWD | DHC | None |  |
| MS | MS-FMINCON-ADJ-LOG | MS-NL2SOL-FWD-LOG | MS-DHC-LOG | – | LOG |
|  | MS-FMINCON-ADJ-LIN | MS-NL2SOL-FWD-LIN | MS-DHC-LIN | – | LIN |
| eSS | eSS-FMINCON-ADJ-LOG | eSS-NL2SOL-FWD-LOG | eSS-DHC-LOG | eSS-NOLOC-LOG | LOG |
|  | eSS-FMINCON-ADJ-LIN | eSS-NL2SOL-FWD-LIN | eSS-DHC-LIN | eSS-NOLOC-LIN | LIN |

- **MS**: multi-start local optimization.
- **eSS**: enhanced scatter search metaheuristic with three different local methods:
- **NL2SOL-FWD**: the nonlinear least-squares algorithm NL2SOL, using forward sensitivity analysis for evaluating the gradients of the residuals. The use of NL2SOL [14] has recently been advocated for parameter estimation by [22]. Additionally, [46] showed that least-squares algorithms with residual sensitivities computed using forward sensitivity analysis outperform many alternative approaches.
- **FMINCON-ADJ**: the interior point algorithm included in FMINCON (MATLAB and Optimization Toolbox Release 2015a, The MathWorks, Inc., Natick, Massachusetts, United States), using adjoint sensitivities for evaluating the gradient of the objective function. This method has been shown to outperform the least-squares method using forward sensitivities for large-scale models [20, 21], due to the accelerated gradient evaluation.
- **DHC**: a gradient-free dynamic hill climbing algorithm. This algorithm has been proposed by [12] and outperformed several alternative approaches in a recent study [58]. In our experience, this method is competitive when the gradient is numerically difficult to evaluate, e.g., if objective function values are corrupted by numerical integration errors.

The considered global strategies and local methods are a representative subset that covers distinct approaches, which have been shown in the past to exhibit competitive performances on a number of problems.

### 2.4 Choice of scaling for the optimization variables

In addition to the optimization methods, we consider two different choices for the scaling of the optimization variables:

- **LIN**: linear scale
- **LOG**: logarithmic scale

While it is possible to consider the model parameters, *p*, directly as optimization variables, several studies suggest that using the logarithms of the model parameters, *q* = log(*p*), improves the performance of local optimization methods [31, 46].

### 2.5 Comparison of optimization methods

The performance of optimization methods can be compared using several evaluation criteria. Ideally, a criterion should be:

1. **single, interpretable quantity**
2. **comparable across models and methods** (to enable an integrated analysis)
3. account for **computation time and objective function value**

A number of evaluation criteria have been used in the literature to compare the performance of optimization methods, e.g., *dispersion plots* of objective function value versus computation time and *waterfall plots* showing the ordered objective function values found by the different searches. These and other plots are reported in the Supplementary Information, Figs. S1–S14. Alternative criteria are *performance profiles* [16] which report for a given set of optimization problems how often one algorithm was faster than all others. The required assumption that all algorithms converge is relaxed in for *data profiles* [41] by considering the decrease in objective function value and reporting the fraction of solved problems as a function of the budget per variable. While all these plots are useful tools, they do not provide a single, interpretable quantity and fail in other ways.

Upon consideration of a variety of different evaluation criteria, we decided to adopt a workflow consisting of several steps, which lead to a newly proposed metric that is a distillation of the information obtained in previous steps. The workflow considers the following criteria:

1. Convergence curves
2. Fixed-budget scenario and fixed-target scenario
3. Dispersion plots of the success rate versus average computation time
4. Overall efficiency (OE)

The first step is to evaluate *convergence curves*, which show the objective function value as a function of computation time (Fig. 1A). For eSS, the convergence curves are constructed from single searches as they reach the predefined maximum CPU time. For MS optimization, each convergence curve corresponds to a sequence of local searches and continues until the predefined maximum CPU time was reached.

**Figure 1:**
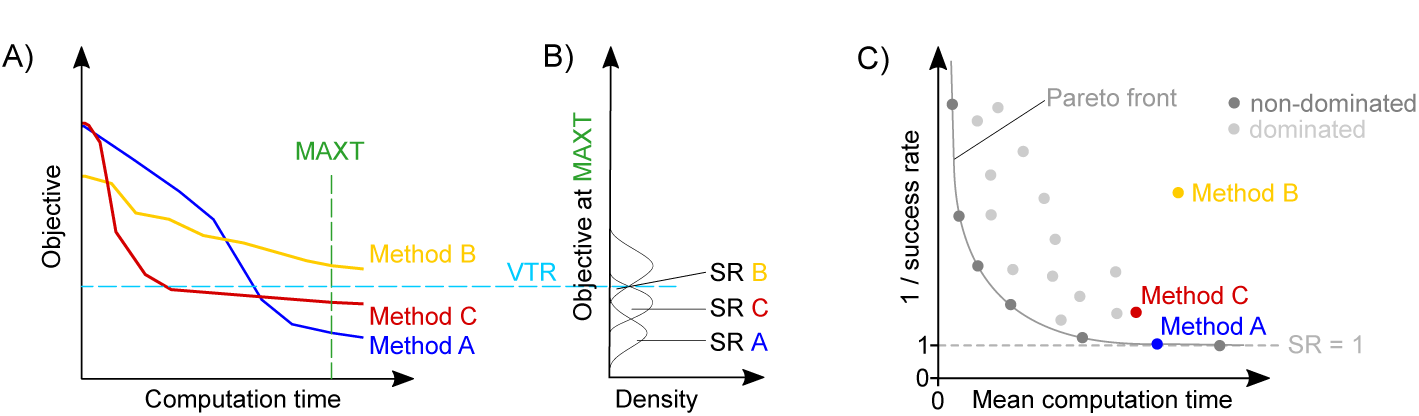
Illustration of performance criteria. A) Convergence curves for three different methods. Shaded areas show the range of all runs, while solid lines represent their median. The dashed horizontal line is the value to reach (VTR), that is the maximum objective function value that can be considered a successful result. The dashed vertical line is the maximum time allowed (MAXT). B) Dispersion plot of objective value after the maximum time allowed and the derived success rates (SR). The SR is the area under the curve where objective ≤ VTR. C) Success rate and computation time. Points indicate individual methods. The Pareto front is the set of non-dominated methods. Methods to the right or above the Pareto front are dominated by other methods with either shorter computation time or higher success rate. Filled areas show the average computation time <*t*>_succ_ required to obtain a successful run for the respective method.

The information encoded in the convergence curves is in the second step summarized by considering a *fixed-budget scenario* and a *fixed-target scenario*, as proposed by [24]. In the fixed-budget scenario, the distribution of the objective function for a given computation time is considered, meaning that a vertical line is drawn. In the fixed-target scenario the distribution of time points is considered at which a desired objective function value or value to reach (VTR) is reached, meaning that a horizontal line is drawn. Once an optimization has reached the desired VTR (horizontal view), it is considered successful. The success rate (SR) of an algorithm is the fraction of searches that reached the VTR within this maximum computation time, MAXT (Fig. 1B). Complementary, we evaluate the average computation time required by an algorithm, <*t*>, which is the minimum of the time required to reach VRT and MAXT. In the third step, we consider *dispersion plots of the success rate versus average computation time* to study the relation of the two quantities (Fig. 1C). Note that this dispersion plot may reveal in some cases a Pareto set structure, consisting of algorithms which provide an optimal trade-off between conflicting goals (in this case, high success rate and low computation time): it is not possible to improve one of its objectives without worsening the other. We are interested in methods that are located towards the bottom (i.e. high success rate) and left (i.e. low computation time) of this plot. Therefore, in the fourth step, we quantify the trade-off between success rate and average computation time using a novel metric called *overall efficiency* (OE). The OE for method *i* on a given problem is defined as:

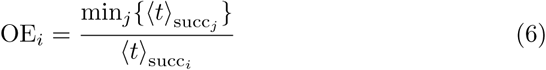

where <*t*>_succ_i__ is the average computation time we need to run method *i* to obtain one successful run. It is calculated as <*t*>_succ_i__ = <*t*>_*i*_/*SR*_*i*_, where <t>_*i*_ and SR_*i*_ are the average computation time and the success rate of method *i* for that problem. The computation time <*t*>_succ_i__ is directly related to the area in the dispersion plot (Fig. 1C); accordingly, the OE is the ratio of the minimal area and the area for a given algorithm. The inverse of the overall efficiency, 1/OE_*i*_, quantifies how much longer one has to run method *i* – compared to the best method – in order to find a good solution. The OE ranges between 0 and 1; for each particular problem the best performing method achieves the maximum score, OE = 1. To evaluate methods on a set of optimization problems, we compute a method’s cumulative overall efficiency as the sum of its OEs for the individual problems. The method with highest cumulative OE will be the one exhibiting the best trade-off between success rate and computation time for the set of problems.

In summary, our workflow considers multiple criteria and summarizes the trade-off between computational complexity and success with the OE. This novel metric fulfils all the afore-defined criteria.

### 2.6 Benchmark problems

In this study, we consider seven benchmark problems based on previously published kinetic models [7, 9, 30, 36, 40, 50, 56] which describe metabolic and signalling pathways of different organisms (from bacteria to human). These problems possess 36 to 383 parameters and 8 to 104 state variables. The data points are collected under up to 16 experimental conditions, corresponding to the number of required numerical simulations. The features of all problems are summarized in Table 2. The benchmarks B2–B5 had been previously included in the BioPreDyn-bench collection [58], and BM1 & BM3 were used in [21].

**Table 2:** Main features of the benchmarks. The model IDs follow the nomenclature in [58] and [21].

| ID | B2 | B3 | B4 | B5 | BM1 | BM3 | TSP |
| --- | --- | --- | --- | --- | --- | --- | --- |
| Original ref. | [7] | [30] | [56] | [36] | [50] | [9] | [40] |
| Organism | *E. coli* | *E. coli* | Chinese hamster | Generic | Mouse | Human | Generic |
| Description | Metabolic | Metabolic | Metabolic | Signaling | Signaling | Signaling | Metabolic |
| level |  | & transcrip. |  |  |  |  |  |
| Parameters | 116 | 178 | 117 | 86 | 383 | 219 | 36 |
| Dynamic states | 18 | 47 | 34 | 26 | 104 | 500 | 8 |
| Observed states | 9 | 47 | 13 | 6 | 12 | 5 | 8 |
| Experiments | 1 | 1 | 1 | 10 | 1 | 4 | 16 |
| Data points | 110 | 7567 | 169 | 960 | 120 | 105 | 336 |
| Data type | measured | simulated | simulated | simulated | measured | measured | simulated |
| Noise level | real<sup>▲</sup> | no noise | variable<sup>◆</sup> | $\sigma = 5\%$<sup>★</sup> | real<sup>▲</sup> | real<sup>▲</sup> | $\sigma = 5\%$<sup>★</sup> |

<sup>▲</sup>Noise levels are unknown as real measurement data are used.
<sup>◆</sup>Noise levels differ for readouts.
<sup>★</sup>Noise levels are proportional to the signal intensity.

### 2.7 Implementation

The benchmark problems have been implemented in MATLAB (MathWorks, Natick, MA, USA) using the AMICI toolbox [20], a free MATLAB interface for SUNDIALS solvers [26]. The optimization methods have been implemented as MATLAB scripts calling solvers from the MATLAB Optimization Toolbox and from the MEIGO toolbox [18], and making use of the efficient gradient computation provided by the the AMICI toolbox. The code necessary for reproducing the results reported here is available at Zenodo https://doi.org/10.5281/zenodo.1160343.

## 3 Results and Discussion

### 3.1 Comprehensive evaluation of the considered optimization methods on the benchmark problems

To assess the performance of the different optimization methods, we solved the 7 benchmark problems using the 14 optimization methods listed in Table 1. The optimization methods were run 10 times, each time until the predefined, maximum problem-specific CPU time (Supplementary Information, Tab. S1) was reached, resulting in an overall computational effort of ~400 CPU days. The convergence curves for all optimization methods on all problems were evaluated (see Fig. 2A for a representative example and Supplementary Information, Figs. S15–S28 for the complete set). Numerical values of the horizontal and vertical views of said curves are provided in Tables S1–S4, and graphically in Figs. S37–S40. As expected, the optimization results indicate that the performance of the optimization methods varies substantially among the benchmark problems. This is in agreement with previous studies [31, 58].

**Figure 2:**
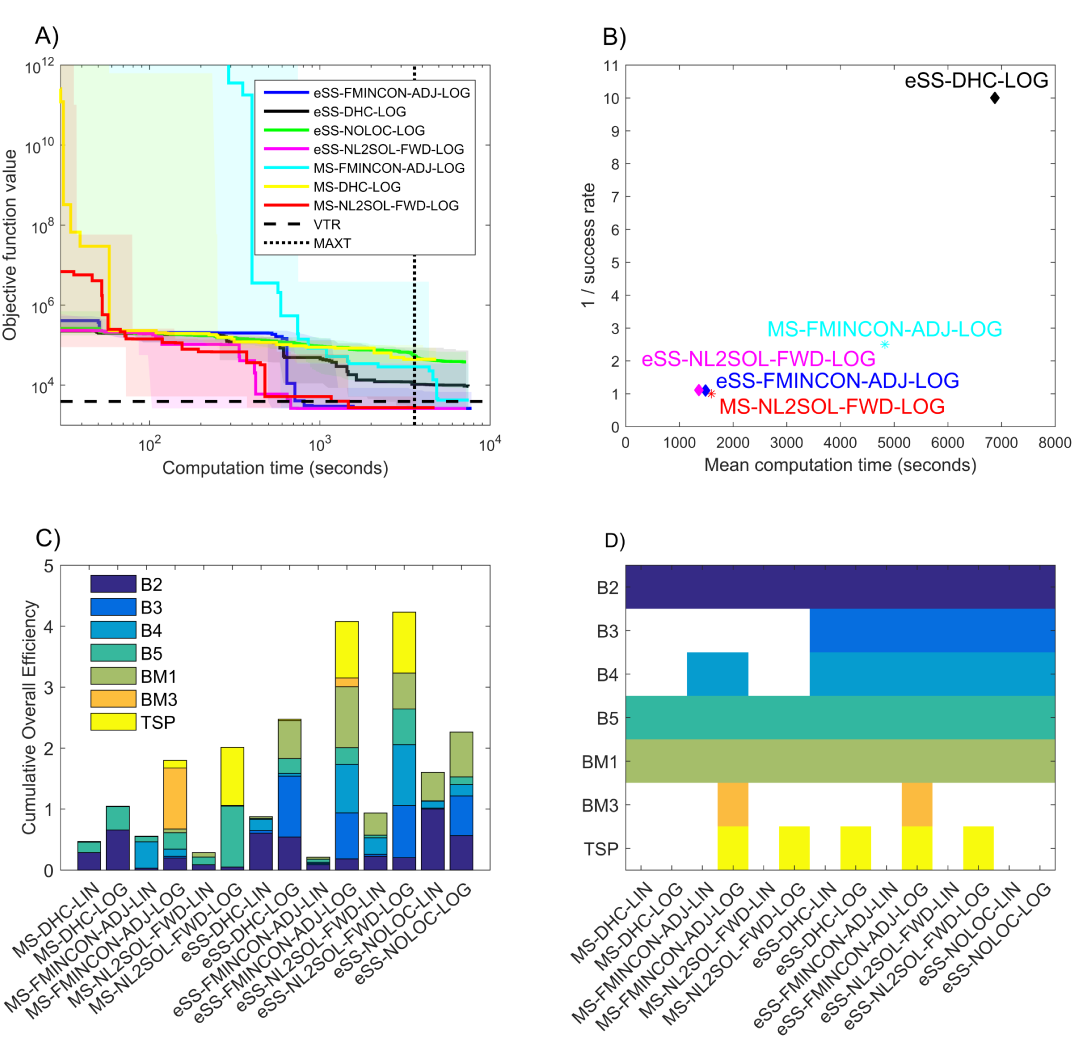
Results of performance evaluation. A) Convergence curves of the different methods for benchmark TSP. Results for the remaining benchmarks are reported in the Supplementary Information. B) Average computation time of each method versus the inverse of its success rate for benchmark TSP. Methods with zero success rate are not shown. Results for the remaining benchmarks are reported in the Supplementary Information. C) Cumulative overall efficiency: Each method is represented by a stack of the OEs observed for the individual benchmark problems. The maximum possible score is the same as the number of benchmarks, i.e. seven. D) Successful methods for each benchmark are shown in colour; methods which never succeeded for a given problem are shown in white.

For the quantitative evaluation, we selected a VTR for each benchmark which provides a solution with a good visual agreement between model output and data. Since the choice of a specific VTR is problematic, we repeated the analyses for four different values, finding that the relative ranking of the methods in terms of performance was robust to changes in the VTR. Hence, in the following subsections, we report results for a reference VTR; results for other choices of VTR including larger and smaller values are shown in the Supplementary Information, Figs. S29–S40.

In the following, we present the key findings of our analysis and address, amongst others, the question of which is the most efficient method for performing parameter optimization. The detailed evaluation results are presented in the Supplementary Information.

### 3.2 Gradient-based local searches outperform gradient-free local searches

Our comprehensive evaluation clearly shows that high-quality sensitivity calculation methods provide a competitive advantage to local methods that exploit them. Optimization using adjoint and forward sensitivity analysis (FMINCONADJ and NL2SOL-FWD) usually outperform the gradient-free alternative (DHC). This is reflected in the dispersion plots (see, e.g., Fig. 2B) and in a higher cumulative OE (Fig. 2C) and holds for MS and eSS settings. Notably, successful optimization of BM3 for the given computational budget required adjoint sensitivity analysis in combination with optimization in the log-scale (Fig. 2D).

### 3.3 Enhanced scatter search outperforms multi-start local optimization

Our results show that MS is usually sufficient to find a good solution, given the same computation time as eSS (Fig. 2D). However, eSS were generally more efficient than MS (Fig. 2C). On average a 2-fold improvement of the OE is observed, almost independent of the local method. The reasons for the efficiency improvement is probably that eSS starts the local searches from promising points found through advanced exploration and recombination strategies. In this regard, it can be considered as an “advanced multi-start” [52].

### 3.4 Optimization in logarithmic scale outperforms optimization in linear scale

Previous studies reported that the transformation of the optimization variable to log-scale improves the reliability and computational efficiency of local methods [31, 46]. Our findings corroborate these results and show for the first time that also global optimization methods are more efficient when using log-scale (LOG) than linear-scale (LIN). Overall, we observe an average improvement of the cumulative OE by a factor of 2 (Fig. 2C). Indeed, for some problems (BM3, TSP), reasonable fits could only be obtained using the log-transformed parameters (Fig. 2D).

### 3.5 Best performing method

The comparison of all methods reveals that eSS-NL2SOL-FWD-LOG possesses the best overall efficiency on the considered benchmark problems and settings, closely followed by eSS-FMINCON-ADJ-LOG (Fig. 2C). The difference in performance between both methods is small; indeed, if different VTRs are chosen eSS-FMINCON-ADJ-LOG can become the best performer (Figs. S33, S34, S36). Complementary, eSS-FMINCON-ADJ-LOG is the only method that successfully solves all problems (Fig. 2D), while the method with the best performer (eSS-NL2SOL-FWD-LOG) fails for BM3, possibly due to the very large number of states and parameters of this problem. In summary, our performance evaluation hence suggests the use of eSS-FMINCON-ADJ-LOG.

## 4 Conclusion

In this paper we have presented a comparative evaluation of state-of-the-art optimization methods for parameter estimation in systems biology. We have applied these methods to a benchmark problems of different sizes (medium to large) and complexities. To compare the different methodologies in detail, we have used a multi-criteria workflow, exploring several possible ways of assessing the performance of optimization methods for this task. We have reported results using a number of selected indicators and evaluation tools. Furthermore, we have introduced the concept of overall efficiency (OE), which quantifies the trade-off between success rate and computation time, providing a numerical indication of the most efficient method. We have found that this metric is a convenient summary of the comparative performance of a method on a set of problems.

A central goal of our work was to re-examine past results regarding the performance of multi-start and metaheuristics (i.e. enhanced scatter search). Firstly, we have confirmed that multi-start local optimization is a powerful approach [27,46] as it solved most considered benchmark problems in a reasonable time. The only exception is B3, a problem for which numerical simulation fails for many parameter points. Secondly, we verified that the enhanced scatter search metaheuristic often possesses higher success rates and efficiency compared to plain multi-start optimization methods [22]. However, the difference of a factor of two was smaller than suggested by several previous studies and will likely depend on the set of benchmark problems. Furthermore, the average improvement by a factor of two is smaller than the variability across benchmarks, implying that for many problems the use of multi-start methods is still beneficial (e.g., BM3). Thirdly, our results confirm that a purely global optimization strategy (i.e. not combined with a local method) is less efficient than a hybrid one. Finally, we have assessed the influence of parameter transformations, concluding that optimizations in logarithmic scale clearly outperform those in linear scale.

We considered two sophisticated gradient-based methods, FMINCON with adjoint sensitivities and NL2SOL with forward sensitivities, whose use was mostly beneficial. A gradient-free local method, DHC, was found to be less precise than the gradient-based counterparts, although its use may still be advantageous in problems with numerical issues that limit the efficacy of gradientbased techniques.

Overall, the best performing method in our tests was eSS-FMINCON-ADJLOG, that is, a hybrid approach combining the global metaheuristic eSS with the local method FMINCON, provided with gradients estimated with adjoint-based sensitivities. This was the only method that succeeded in calibrating all the benchmarks and it also achieved a good overall efficiency. To facilitate the application of this and other methods, we provide their implementations in the Supplementary Material. In the case of the best performing method, our solver is – to the best of our knowledge – the first publicly available implementation. Accordingly, our study provides access to a novel optimizer applicable to a broad range of application problems in systems biology.

